# The Crux toolkit for analysis of bottom-up tandem mass spectrometry proteomics data

**DOI:** 10.1101/2022.10.02.510538

**Authors:** Attila Kertesz-Farkas, Frank Lawrence Nii Adoquaye Acquaye, Kishankumar Bhimani, Jimmy K. Eng, William E. Fondrie, Charles Grant, Michael R. Hoopmann, Andy Lin, Yang Y. Lu, Robert L. Moritz, Michael J. MacCoss, William Stafford Noble

## Abstract

The Crux tandem mass spectrometry data analysis toolkit provides a collection of algorithms for analyzing bottom-up proteomics tandem mass spectrometry data. Many publications have described various individual components of Crux, but a comprehensive summary has not been published since 2014. The goal of this work is to summarize the functionality of Crux, focusing on developments since 2014. We begin with empirical results demonstrating our recently implemented speedups to the Tide search engine. Other new features include a new score function in Tide, two new confidence estimation procedures, as well as three new tools: Param-medic for estimating search parameters directly from mass spectrometry data, Kojak for searching cross-linked mass spectra, and DIAmeter for searching data independent acquisition data against a sequence database.

## 1 Introduction

Continual technological advances in mass spectrometry instrumentation, which yield higher throughput, increased data depth, accuracy and precision, and innovative orthogonal modes of ion measurement require concomitant advances in analytical methods. Crux is an open source software project that implements a variety of state-of-the-art algorithms for interpreting bottom-up tandem mass spectrometry proteomics data. The algorithms implemented in Crux are described in 40 scientific papers, cited a total of 6,413 times and with an H-index of 25.^1^ A typical Crux user is unlikely to read this large corpus of papers; hence, the goal of this paper is to provide an overview of Crux, with a focus on developments that have been introduced since our last overview paper in 2014 [1].

The field of computational mass spectrometry is broad, and Crux necessarily occupies a particular niche within that field. In particular, Crux focuses primarily on the initial stages of tandem mass spectrometry analysis: the assignment of peptides to spectra, with associated measures of statistical confidence at the level of spectra, peptides and proteins. Crux includes four database search tools, two for standard search (Tide and Comet), one for searching against a database of cross-linked peptides (Kojak), and one for searching data-independent acquisition (DIA) data (DIAmeter) (Figure 1). Also included is the Bullseye tool for assigning high-resolution precursor masses to MS2 spectra, a machine learning post-processor (Percolator), a separate tool for assigning confidence estimates to various types of discoveries (assign-confidence), and a spectral counting tool (spectral-counts). Practically speaking, Crux is a command line tool, written in C++. Source code is available, and we also provide pre-compiled binaries for use on Microsoft Windows, MacOS and Linux operating systems from http://crux.ms.

**Figure 1.**
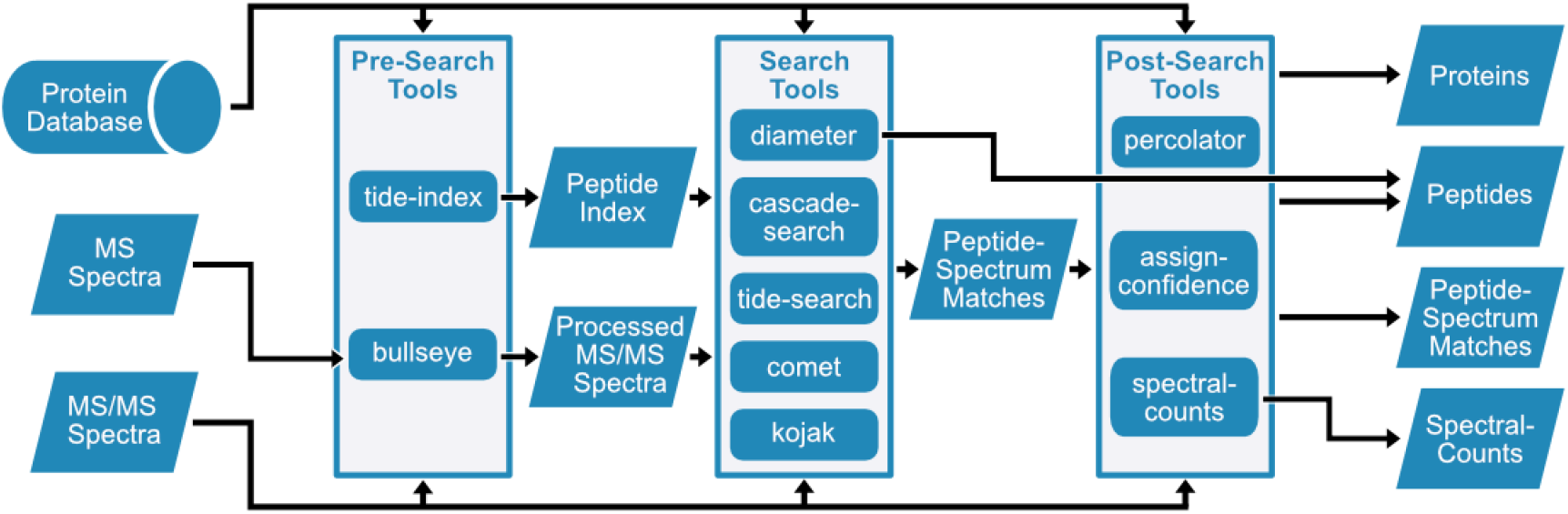
Overview of tools in Crux. Bullseye assigns high resolution precursor m/z values to tandem mass spectra. Crux includes two DDA search tools, Tide and Comet, plus a variant of Tide called cascade-search, described in Section 3.2. DIAmeter searches data-independent acquisition data, and Kojak searches cross-linked mass spectra. Percolator is a machine learning post-processor, assign-confidence estimates statistical confidence estimates directly from search results, and spectral-counts computes several types of protein abundance measures using spectral counting.

In this paper, we provide an overview of new features in Crux, beginning with empirical results demonstrating our recently implemented speedups to the Tide search engine. Other new features include a variety of new score functions in Tide, several enhancements to the Comet search engine, two new confidence estimation procedures, as well as three new tools: Param-medic [2, 3], Kojak [4], and DIAmeter [5].

## 2 Methods

### 2.1 datasets

For the benchmarking in Section 3.1–3.2, we selected at random one raw file (20190601_QX6_JoMu_ SA_uPac200cm_HepG2_f4.raw) from a human sample in a recent large-scale study [6] (PRIDE accession PXD014877). The file contains 178,024 spectra. For the Param-Medic analyses in Section 3.4.1 we analyzed all 26 RAW files associated with PRIDE project PXD004424.

Crux is capable of analyzing RAW files directly, but only on a Windows machine. Because our analyses were performed on Linux systems, all RAW files were first converted to an open format using ThermoRaw-FileParser [7].

### 2.2 Protein databases

Searches were conducted against the human reference proteome file (uniprot-proteome_UP000005640.fasta) downloaded from Uniprot on Feb 3, 2022. The fasta file contains canonical and isoform protein sequences.

### 2.3 Search engines

In the comparison of search engines, we tried to ensure that comparable settings were employed between Comet and Tide (Table 1). Note that when switching to the exact p-value score function in Tide, we were obliged to set `mz-bin-width to 1.0005079`, and for the combined p-value score function, we used `-mz-bin-width` 1.0005079 and `--fragment-tolerance 0.02`. The database search was carried out on a Linux server equipped with an Intel Xeon CPU E5-2640 v4 2.40GHz processor with 20 cores and 1TB SSD storage. Although both Comet and TIde allow multiple threads, the searches performed here use a single thread.

**Table 1:**
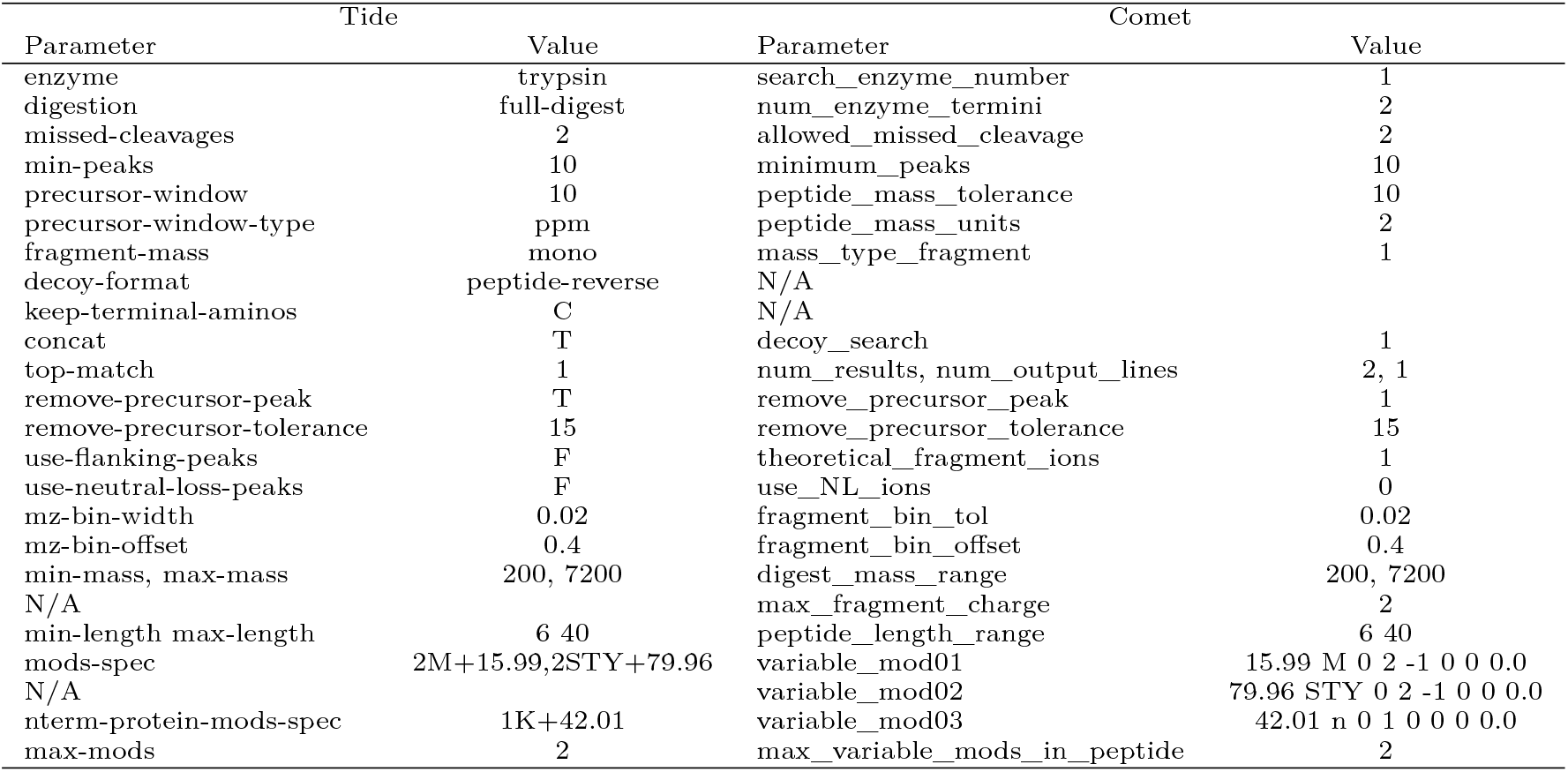
Parameter settings for Comet and Tide.

**Table 2:** Running time comparison of two versions of Tide. The table shows the running time in seconds of Tide with four different score functions (XCorr, Tailor, exact p-value and combined p-value) and Comet in the old (v3.2) versus the new version (v4.1-36) of Crux. The search was performed with data described in Sections 2.1–2.3.

| Search | Old crux | New crux |
| --- | --- | --- |
| Tide XCorr | 1,230 | 365 |
| Tide Tailor | 1,250 | 284 |
| Tide p-value | 3,640 | 813 |
| Tide combined | 16,300 | 6,910 |
| Comet | 1,140 | 1,670 |

## 3 Results

### 3.1 Tide speedups and new score functions

We begin our analysis with a timing comparison of various score functions, as implemented in Crux’s two DDA database search tools, Tide and Comet. In its initial implementation, Tide was markedly faster than competing search engines [8]. However, subsequent modifications to the code to implement new features and new score functions led to a decrease in Tide’s efficiency. Consequently, we recently overhauled the Tide code with a focus on speeding it up, yielding a three-fold increase in speed relative to the previous version of Tide (Table 3.1). As a result, Tide is now quite efficient (Figure 2A), capable of searching the tryptic human proteome at *∼*750 spectra/s. In particular, in its fastest mode, Tide searching is around 4.5 times faster than Comet searching.

**Table 3:** Running time comparison for Tide and Comet. All times are reported in seconds. The data corresponds to Figure 2A.

| Number of Peptides | Tide XCorr | Tide Tailor | Tide p-value | Tide combined | Comet E-value |
| --- | --- | --- | --- | --- | --- |
| 60,896,400 | 241 | 251 | 646 | 7,140 | 1,100 |
| 40,484,062 | 211 | 219 | 581 | 5,570 | 807 |
| 24,777,903 | 188 | 193 | 566 | 4,450 | 694 |
| 13,635,673 | 171 | 173 | 551 | 3,660 | 631 |
| 7,461,453 | 159 | 161 | 531 | 3,070 | 599 |

**Figure 2.**
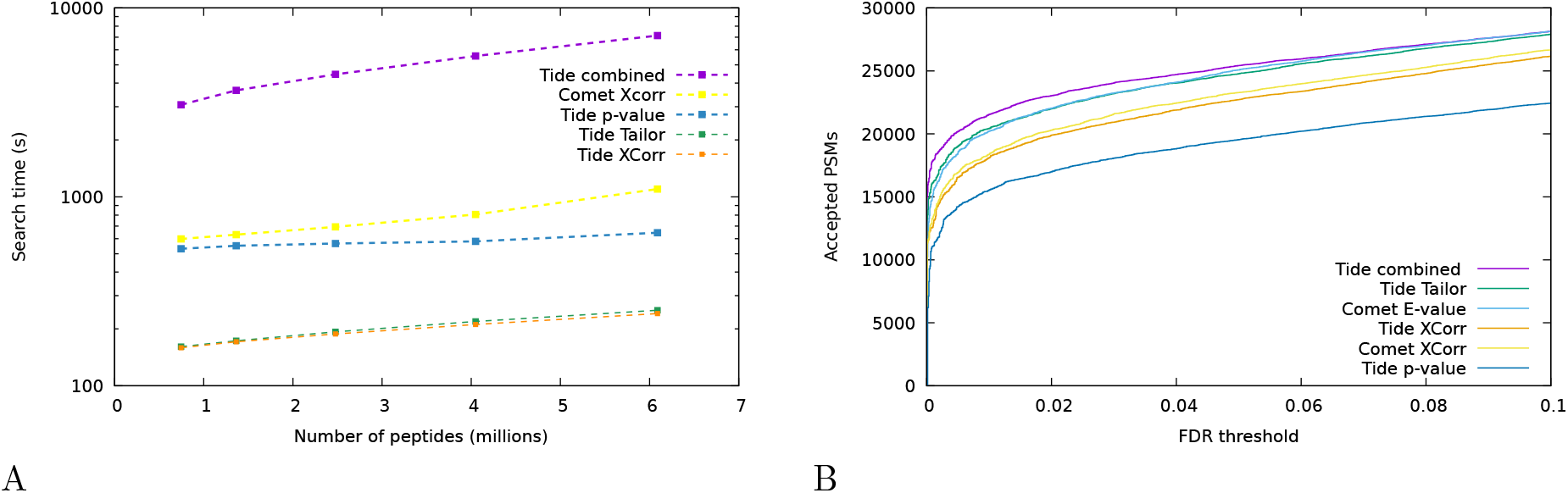
Comparisons of search tools. (A) The figure plots the total running time of Tide and Comet, as a function of database size. The series correspond to Comet and Tide with four different score functions (XCorr, Tailor, exact p-value and combined p-value). The search was performed with data described in Sections 2.1–2.3. The proteome was randomly downsampled to contain the specified number of peptides. Detailed timing information is provided in Table 3.1 (B) The figure plots the number of accepted PSMs as a function of q-value threshold. The series correspond to Comet (with and without a pre-computed peptide index) and Tide with four different score functions (XCorr, Tailor, exact p-value and combined p-value). The search was performed with data described in Sections 2.1–2.3. All q-values are assigned using target-decoy competition, as implemented in `assign-confidence` in Crux.

Tide recently introduced a new scoring scheme, called Tailor calibration, which calibrates the top PSM score relative to the full distribution of scores were generated during the database search. In this sense, it is similar to the E-value calibration implemented in Comet [9]. Specifically, Tailor considers the PSM scores *s*_1_, *s*_2_, …, *s*_*N*_, (in decreasing order) when matching one experimental spectrum to a set of *N* candidate peptides. Tailor calibration identifies the 99th quantile of this distribution by selecting the PSM score at the position *i*^*\**^ = [*N/*100], where [.] denotes the standard rounding operation. The Tailor method calibrates the top PSM score *s*_1_ by 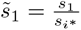. Tailor is thus a simple and quick method for score calibration.

From the user’s perspective speed is only useful in conjunction with accurate results. Accordingly, we compared the statistical power of various search strategies by counting the number of peptide-spectrum matches (PSMs) accepted at a 1% false discovery rate (FDR) threshold, as estimated using target-decoy competition. The results show several expected trends (Figure 2B). First, the raw XCorr score, as implemented in either Comet or Tide, does not perform as well as the corresponding calibrated score (the Comet E-value or Tide’s Tailor score [10]). Tide also includes an alternative calibrated score, the “exact p-value,” that is estimated using a dynamic programming procedure [11]. However, the exact p-value is designed to work with data that is generated using low-resolution precursor scans, so it actually yields decreased statistical power on the high-resolution data we used. Tide’s “combined p-value” score is designed to combat this problem by combining the exact p-value with another dynamic programming procedure that operates on pairs of amino acids [12]. This score yields the best overall performance but is markedly slower to compute.

### 3.2 Confidence estimation procedures

The Tide search engine now supports two new procedures to improve statistical confidence estimation. The first procedure, known as cascade search [13], aims to boost statistical power—i.e., the number of peptides detected at a specified FDR threshold. Cascade search is applicable when the peptide database can be divided into groups *a priori*, and the groups can be ordered from more likely peptides toward more rare peptides. Cascade search works by sequestering at each stage any spectrum that is identified with a specified statistical confidence and then searching the remaining spectra against the next database in the list. For instance, such a cascade of databases could include fully tryptic, semitryptic, and nonenzymatic peptides or peptides with increasing numbers of modifications.

To demonstrate the empirical benefit of cascade search on a sample dataset, we analyzed a sample dataset in two ways: using a single peptide database followed by FDR control with target-decoy competition (TDC), and using cascade search with respect to a series of databases created using fully tryptic, semitryptic and non-enzymatic digestion. In Crux, cascade search is implemented as a separate command (cascade-search) that takes as input one or more spectrum files plus a comma-separated list of Tide indices. For this experiment, we used the same human dataset as before (described in Sections 2.1–2.3). We observe that at 1% FDR, cascade search accepts 27,400 PSMs, whereas a single-database Tide search accepts only 20,448, 25,325, or 23,046 PSMs, depending on whether the database is tryptic, semitryptic or non-enzymatic. Thus, cascade-search leads to an increase in the number of accepted PSMs between 8–34% at 1% FDR.

Note that the cascade search procedure, in this case, is somewhat inefficient because the three databases are supersets of one another; e.g., all tryptic peptides are also included in the semitryptic database. To avoid this inefficiency, Crux provides an auxiliary command, subtract-index, that will remove from one Tide index all peptides that occur in a second index.

The second new procedure aims to reduce the variance in FDR estimates that is intrinsic to any decoy-based confidence estimation method. The procedure, called “average target-decoy competition” (aTDC) [14, 15], works by searching a given set of spectra against a collection of peptide databases: one database containing target peptides and multiple database containing shuffled decoy peptides. In Crux, aTDC is implemented via the `num-decoys-per-target`. Setting this parameter to any integer *>* 1 will cause Tide to carry out aTDC.

We demonstrated the utility of aTDC using the same human dataset as before (described in Sections 2.1–2.3). In practice, averaging is most useful when the total number of discoveries is small, because in this setting the decoy-induced variance in the estimated FDR can have a substantial impact on the results. Accordingly, to simulate such a scenario, we searched a database containing 100 proteins selected at random from the human proteome. In this setting, the variability that we observe in the FDR estimates from standard TDC is substantially reduced when we use aTDC with five decoys per target (Figure 3). For example, at a 1% FDR threshold, the standard deviation in the number of accepted PSMs decreases by 83%, from 42 to 7.

**Figure 3.**
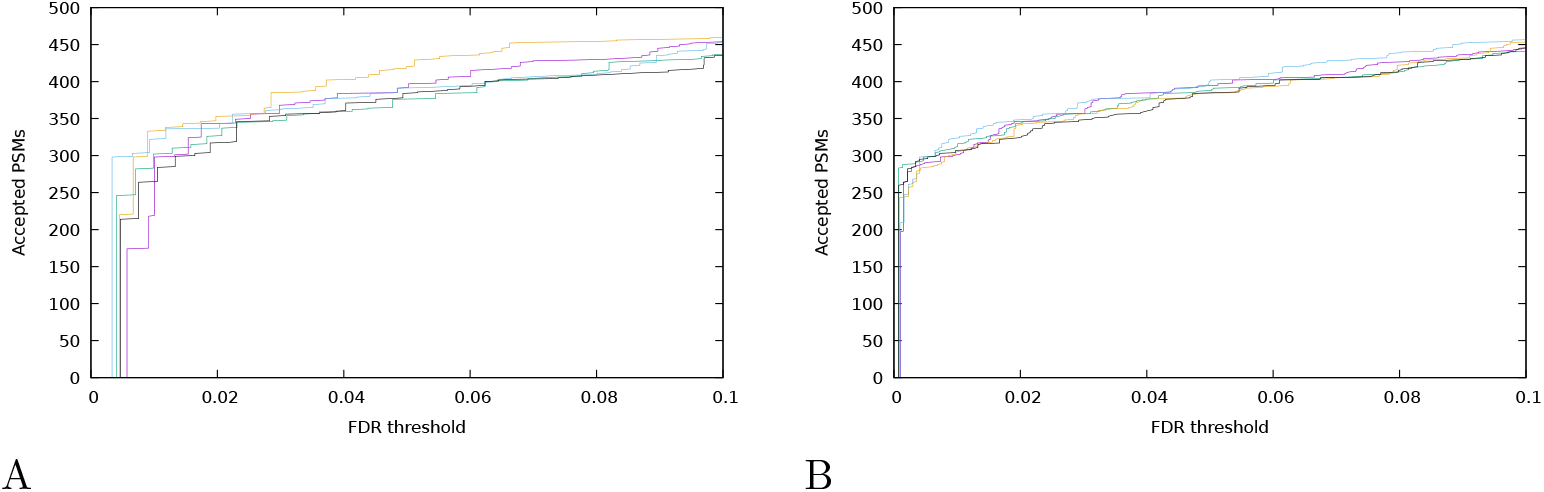
Average target-decoy competition reduces decoy-induced variance. (A) The figure plots the number of accepted PSMs (y-axis) as a function of FDR threshold (x-axis), for searches against databases of varying size. Each series is generated by searching a different, randomly shuffled decoy database. (B) Similar to panels (A), except that each of the five series in the plot corresponds to FDR estimates from aTDC, using five decoys per target.

### 3.3 Comet updates

Since the last Crux overview paper in 2014, the Comet search tool has incorporated many updates and bug fixes.

One feature that has been extended for analysis flexibility, based on requests by various researchers attempting to optimize specific analysis, is the control of how variable modifications are applied. This includes distance constraints of modifications from peptide or protein termini, forcing the requirement of a modification to be present in a peptide, including the ability to specify the minimum and maximum number of each variable modification, controlling whether or not a variable modification can appear on the C-terminal residue, and consideration of neutral loss peaks on those fragment ions that contain a variable modification.

Comet was also one of the first search tools to support the Proteomics Standards Initiative’s Extended Fasta Format (PEFF) [16]. Comet’s initial published PEFF support included the ability to search PEFF database files to analyze the annotated modifications and single amino acid substitutions [17]. More recently, Comet’s PEFF support has been extended to include the ability to analyze “VariantComplex” annotations which encode sequence variations that are more complex than a single amino acid substitution. VariantComplex annotations can encode deletions, insertions, and combinations of the two, which allows the PEFF database to encapsulate sequence variations such as protein isoforms within a single sequence entry.

Comet was also extended to support the real-time search application that was initially implemented in the Schweppe lab’s Orbiter platform for real-time instrument control [18]. Subsequently, Comet’s real-time search application has been adopted by Thermo Scientific and is now available for real-time analysis on their Tribrid mass spectrometers, typically for support of tandem mass tag workflows to increase unique data depth

### 3.4 New tools

#### 3.4.1 Param-Medic

The Param-Medic command automatically infers several key characteristics—precursor window size, fragment ion tolerance, and the presence of several common types of post-translational modifications—of a given MS/MS dataset by examining the MS1 and MS2 spectra. The primary goal is to facilitate automated processing of public datasets, when metadata such as instrument settings may be hard to come by. Param-Medic can also be useful to identify problems with a dataset, for example, when the nominal mass accuracy of the data disagrees with the mass accuracy inferred by the program.

To demonstrate Param-Medic’s utility, we downloaded all 26 RAW files associated with PRIDE identifier PXD004424 and subjected them to Param-Medic analysis. Notably, the results suggested a fairly broad range of precursor window sizes, ranging from 16.79 ppm up to 68.48 ppm, whereas the authors of the original study used a 20 ppm window for all of the analyses [19]. To follow up on this assessment, we selected two specific RAW files, one with the minimum inferred window size of 16.79 ppm (151009_exo3_5) and one with the maximum inferred window size of 68.48 ppm (151218_exo4_4). The relationship between the search engine score and delta mass shows a notably broader distribution for the second file, including a handful of outlier points with high Tailor scores (Figure 4), potentially indicative of problematic acquisition.

**Figure 4.**
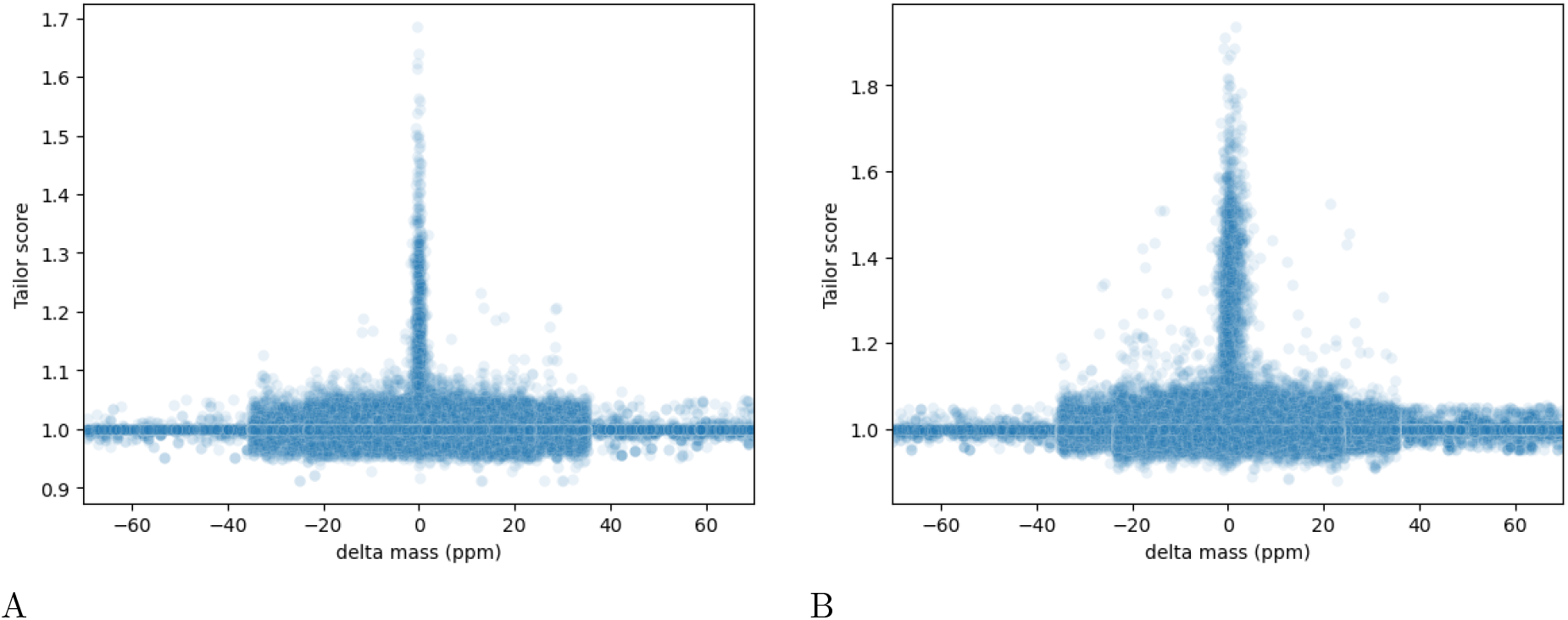
Comparison of precursor acquisition in two different runs. (A) The figure plots, for each PSM produced by searching sample sample 151009_exo3_5 against the human proteome, the Tailor score (y-axis) as a function of the difference between the observed precursor mass and the peptide mass (x-axis). To show a broad range of values, the search was performed with a precursor window size of 70 ppm. For this data, Param-Medic infers a precursor window size of 16.79 ppm. (B) Same as panel (A), but for 151218_exo4_4. The inferred precursor window size is 68.48 ppm.

Note that Param-Medic can be called automatically from within Tide or Comet by using the `auto-modifications-spectra, auto-precursor-window`, and `auto-bin-width options`.

#### 3.4.2 Kojak

Kojak performs database search on mass spectra from cross-linked samples [4]. Here, we describe how to run Kojak on a cross-linked sample from PRIDE project PXD014337 [20] and upload the results into the web-based platform ProXL [21] for visualization. Kojak takes as input mzML spectra data files and a fasta protein sequence file. For this analysis, we analyze the three DSS-linked replicate files using the Cas9_plus10.fasta sequence file (supplemental information). Spectral peaks should be transformed to centroid representation during the conversion from raw spectra to mzML format. Then, it is necessary to tailor a few Kojak parameters to the data:

~~~
fragment_bin_offset = 0.0
fragment_bin_size = 0.01
decoy_filter = DECOY 1
max_miscleavages = 2
min_spectrum_peaks = 25
spectrum_processing = true
top_count = 5
min_peptide_score = 0.25
~~~

These parameters can be specified on the command line or in the Crux parameter file. To run the Kojak analysis on all three data files at once, execute the following command:

~~~
crux kojak --parameter-file kojak.params.txt *.mzML Cas9_plus10.fasta
~~~

This analysis produces a series of files containing cross-linked spectrum matches (CSMs). The files contain the suggested peptide or peptides matched to each spectrum, but these matches must then be validated using a target-decoy approach with Percolator. CSMs are divided into several categories, and we want to validate the intra-protein and inter-protein CSMs. To do this, rename the .txt extensions for *.perc.intra.* *.perc.inter.* files to .pin (e.g. XLpeplib_Beveridge_QEx-HFX_DSS_R1.perc.intra.txt becomes XLpeplib_Beveridge_QEx-HFX_DSS_R1.perc.intra.pin) so that Percolator can read them. Then execute the following command:

~~~
crux percolator --only-psms T --tdc T *.pin
~~~

This command will combine all the Kojak intra-protein and inter-protein CSMs into a single set for Percolator analysis and produce estimated error rates at the CSM-level. Using a q-value threshold of 0.01 to estimate a 1% error rate, 1944 CSMs are returned. Because we know the ground truth in this dataset, we can compare the CSMs to the set of correct results, and find that 1919 are correct, and 25 are incorrect, for an error rate of 1.3%, or approximately the estimated error rate at the chosen threshold.

Visualization of the spectra and CSM annotations is done with ProXL.

#### 3.4.3 DIAmeter

DIAmeter is a library-free database search tool for DIA data [5]. The diameter command in Crux takes as input the DIA data and a user-specified database of proteins, which must first be indexed by the tide-index command. DIAmeter computes a series of scores for each candidate peptide and then calls Percolator internally to produce a ranked list of peptides, with associated confidence estimates (q-values).

A comparative evaluation of DIAmeter appears in the original publication describing the method [5]. Here, we demonstrate how to run the software and show that it gives consistent results on several DIA runs from a recently published study. In this analysis, we use data from a large-scale Alzheimer’s study [22], selecting three runs at random from the hippocampus brain region, batch 1. To search a file “HZR03.mzml” against the Uniprot human proteome (“human.fa”) requires two steps:

1. Create a Tide index from the human reference proteome using the command `crux tide-index human.fa human`.
2. Search the mzML file against the index using the command `diameter --diameter-instrument orbitrap HZR03.mzml human`

For this particular file, DIAmeter detects 12,037 peptides. We also analyzed files from two other samples (HZR07 and HZR10) and detected similar numbers of peptides, with *>*8000 peptides detected in all three runs (Figure 5).

**Figure 5.**
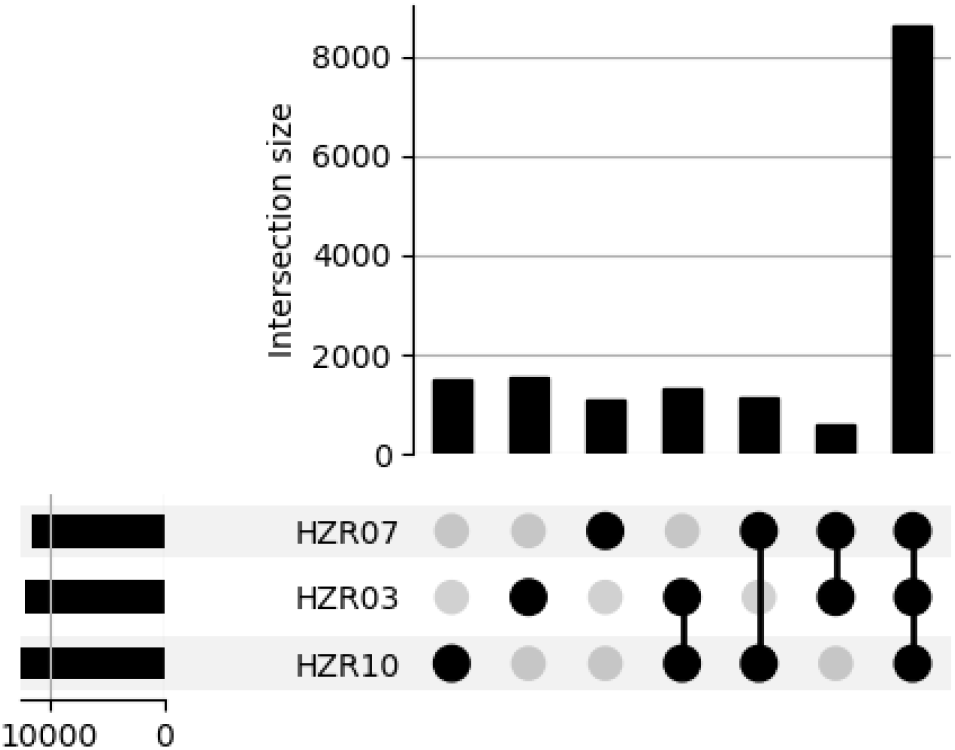
DIAmeter analysis of three Alzheimer’s samples. Three samples from a recent Alzheimer’s study [22] were searched against the Uniprot human reference proteome. The figure shows the number of peptides that were detected at a 1% FDR threshold in all three runs, in any combination of two runs, and in single runs.

## 4 Discussion

Crux provides a rich set of software tools for analyzing proteomics mass spectrometry data. In this paper, we have emphasized the newer aspects of the toolkit, focusing on improvements to our two standard DDA search engines, Comet and Tide, as well as the introduction of several new tools, Kojak, Param-Medic and DIAmeter. Our aim is to ensure that the Crux software can be easily applied to many standard workflows, while also producing accurate results with high statistical power.

As mass spectrometry instrumentation and data collection technology advances, so too do the software tools used to make sense of mass spectrometry data. Accordingly, Crux is under constant development as we work with collaborators and other users of the software to ensure that it addresses their needs. We have a variety of tools planned for future releases, including labeled and label-free quantification tools akin to Libra [23] and FlashLFQ [24], respectively, as well as a mass calibration tool similar to the procedures in MetaMorpheus [25] or MSFragger [26]. Crux users who have specific needs—including new tools to suggest, desired new functionality, or bugs to report—are encouraged to submit an issue to our Github issue tracker, which is linked from the main Crux web page, http://crux.ms.

## Acknowlegment

This work was funded in part by National Institutes of Health grants from the National institute General Medical Sciences R01GM087221, the National Heart, Lung, and Blood Institute R01HL133135, the Office of the Director S10OD026936, and the National Institute on Aging U19AG023122, and by the National Science Foundation award 1920268.

## Table of contents figure

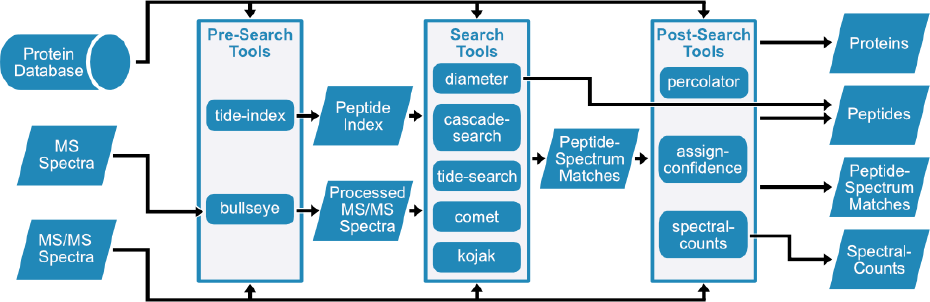

## Footnotes

1 https://scholar.google.com/citations?hl=en&user=Rw9S1HIAAAAJ, Sep 26, 2022

